## Supplementary material for "Transcriptomic profiling of tissue environments critical for post-embryonic patterning and morphogenesis of zebrafish skin": Aman_Saunders_Supplementary_File_1

**Published markers of skin and skin-associated cell types**

| **Cell type** | **Marker (citation)** | **Note** |
| --- | --- | --- |
| Basal Cell | *tp63* (Pellegrini et al., 2001), *apoeb* (Grehan et al., 2001) |  |
| Endothelial Cell | *fgd5a* (Cheng et al., 2012) |  |
| Goblet cells | *muc5*.1 (Okuda et al., 2019) (Jevtov et al., 2014) |  |
| Hypodermis | *col6a3* (Gara et al., 2011), *csf1b* (Lang et al., 2009) |  |
| Ionocytes | *kcnd2* (Pan et al., 2022) |  |
| Iridophore | *gpnmb* (Saunders et al., 2019), *pnp4a* (Lang et al., 2009) |  |
| Leukocyte | *ptprc* (CD45) (Antignano et al., 2019) |  |
| Melanophore | *tyrp1b* (Orlow et al., 1993) |  |
| MLC | *adgrg11* (Alemany et al., 2018; Lin et al., 2019) |  |
| Muscle | *neb* (Labeit et al., 2011) |  |
| NaR ionocytes | *atp1b1b* (Lin et al., 2006) |  |
| Pre-SFC | *col6a3* (general dermal marker) (Gara et al., 2011), *runx2b* (Li et al., 2009) | Simplest diagnosis: *col6a3*+, *runx2b*+; *sp7*-. |
| Periderm | *krt4* (Chen et al., 2011) |  |
| pLL hair cell | *myo7aa* (Gibson et al., 1995) |  |
| pLL mantle cell | *fat1b* (Steiner et al., 2014) |  |
| pLL support cell | *slc1a3a* (Lush et al., 2019) |  |
| PN/Schwann cell | *mbpa* (Takahashi et al., 1985) |  |
| Dermal mesenchyme/  Reticulate dermis | *col6a3* (general dermal marker) (Gara et al., 2011), *postnb* (Fibroblast marker) (Crawford et al., 2015) | Simplest diagnosis: *col6a3*+; *postnb*+; *csf1b*- |
| SFC | *sp7* (Aman et al., 2018) |  |
| Subrabasal cell | *cldna* (Hou et al., 2020) |  |
| Xanthophore | *pax7b* (Nord et al., 2016) |  |
